# SODA: Software to Support the Curation and Sharing of FAIR Autonomic Nervous System Data

**DOI:** 10.1101/2023.10.13.562300

**Authors:** Christopher Marroquin, Jacob Clark, Dorian Portillo, Sanjay Soundarajan, Bhavesh Patel

**Affiliations:** FAIR Data Innovations Hub, California Medical Innovations, Institute, San Diego, CA, USA

## Abstract

Since 2014, the National Institutes of Health (NIH)’s Stimulating Peripheral Activity to Relieve Conditions (SPARC) Program has been supporting research and development of therapeutic devices that modulate electrical activity in the autonomic nervous system (ANS) to improve organ function, also known as bioelectronic medicine. To optimize the reusability of data resulting from ANS-related research, the SPARC Program also supported the development of guidelines for curating and sharing data in line with the FAIR (Findable, Accessible, Interoperable, Reusable) Principles. These guidelines are exhaustive to maximize FAIRness of data but as a result, they are difficult and time-consuming for researchers to implement. To address these challenges, we developed SODA (Software to Organize Data Automatically), an open source and free cross-platform desktop software that guides researchers step-by-step in preparing and sharing their ANS-related data according to the SPARC guidelines. SODA combines intuitive user interfaces with automation to streamline the process and reduce researchers’ time, effort, and error in making their data FAIR. We provide in this paper an overview of SODA and results of testing its performance. We also provide an overview of the impact of SODA which is, to our knowledge, the first researcher-oriented tool for making data FAIR.

## Introduction

The National Institutes of Health (NIH)’s Stimulating Peripheral Activity to Relieve Conditions (SPARC) Program was established in 2014 with the goal of accelerating the development of therapeutic devices that modulate electrical activity in the ANS to improve organ function, also known as bioelectronic medicine (Cracchiolo et al. 2021). Research on bioelectronic medicine had shown tremendous potential to address diverse conditions such as hypertension, heart failure, and gastrointestinal disorders but required further investigation and development for clinical translation (Birmingham et al. 2014; Olofsson 2014), which SPARC aimed to support. In addition, the SPARC Program also supported the development of guidelines for making data resulting from such research optimally reusable in line with the FAIR (Findable, Accessible, Interoperable, Reusable) Principles (Wilkinson et al. 2016). This includes microscopy imaging, electronic measurement data, and more. The goal was to provide a standard way for researchers to organize and share their ANS-related data to facilitate secondary data reuse, enable joint analysis with different datasets, and accelerate discoveries (Quey et al. 2021; Soundarajan et al. 2022). Accordingly, the SPARC data curation and sharing guidelines were developed. The guidelines prescribe the data to be organized according to the standard SPARC Data Structure (SDS) (Bandrowski et al. 2021) and shared publicly on sparc.science, the data portal of the SPARC program (Osanlouy et al. 2021). The SPARC data curation and sharing guidelines have been imposed on all data resulting from SPARC-funded projects since 2017. Since 2022, sparc.science has become an open repository so anyone with ANS-related data can follow the SPARC guidelines to have their dataset published on the SPARC portal. The guidelines are very exhaustive to maximize FAIRness of data but as a result, they are difficult and time-consuming for researchers to implement. While being part of a SPARC-funded research project in 2017, we experienced these challenges firsthand. The process to implement the SPARC guidelines involves several data manipulations to comply with the SDS, which includes organizing data according to a strict folder structure, following file and folder naming conventions, preparing several metadata files, using standard formats for certain data types, and so on. Then, the resulting dataset needs to be uploaded on Pennsieve, the data management platform of the SPARC Program. There, the dataset is reviewed by a team of human curators and a back-and-forth conversation could ensue to address any SDS-compliance issues before the dataset is published on sparc.science. To simplify this process for the data resulting from our SPARC-funded project, we developed Python scripts to automate some of the steps. We presented this automation approach during a SPARC-organized Hackathon in 2018, where it received wide appreciation from other SPARC-funded researchers who were experiencing similar challenges. The project won the Public’s Choice Award at the Hackathon and subsequently received funding from the SPARC Program starting in 2019 for further development of this idea into a desktop software application that anyone can use without coding knowledge. This gave birth to SODA, the software presented in this paper (Patel et al. 2020).

## Method

### Software architecture

SODA is a cross-platform desktop software application developed using Electron, an open-source framework for creating desktop applications using web technologies. Flask is used in the backend of the software to integrate with existing tools that help with complying with the SPARC guidelines, which are mostly developed in Python. Further information about the technical development of SODA can be found in its GitHub repository (Patel, Soundarajan, Marroquin, et al. 2023a). SODA is designed to guide researchers step-by-step through all the requirements for preparing and sharing their data according to the SPARC guidelines. The software combines intuitive user interfaces with automation to streamline the process. A screenshot of the home page of SODA is provided in Figure 1. The development was started in 2019 and is still ongoing as support for more elements of the SPARC guidelines along with additional automation are progressively being implemented. SODA integrates with other SPARC resources such as the SDS validator (Gillespie 2023), the Pennsieve API, and more integrations are planned such that SODA becomes a one-stop tool for anyone wanting to make their ANS-related data FAIR through the SDS and the SPARC data curation and sharing guidelines. A full overview of what SODA supports is available in its user documentation.

**Figure 1:**
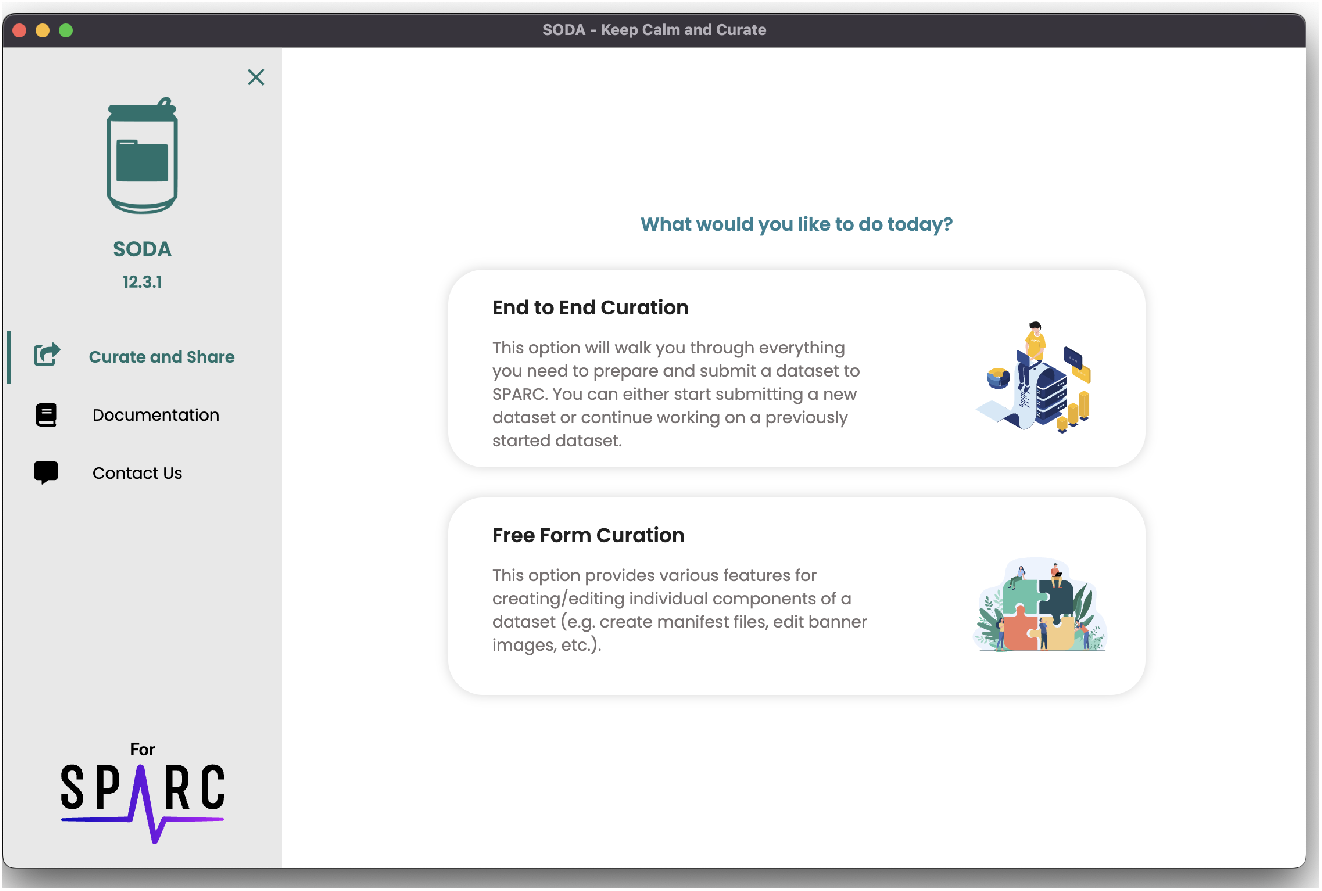
Screenshot of the user interface of the home page of SODA.

### Beta testing

Beta testing was conducted in 2020 after about one year of developing SODA to evaluate its performance. The beta testers consisted of 10 SPARC-funded researchers from 10 different research groups who had never used SODA before but some of them had prepared and submitted data according to the SPARC guidelines without SODA. They were provided a sample dataset and asked to prepare and submit it as per the SPARC guidelines without SODA (Task A) and with only SODA (Task B). For both tasks, they were asked to complete only the steps of the SPARC guidelines supported by SODA at that time (implementing the SDS folder structure, preparing certain metadata files, uploading to Pennsieve, etc.). Half of them (randomly selected) were asked to complete Task A first then Task B while the other half were asked to complete the tasks the other way around. They were requested to report back the time required to complete each task and score out of 5 (1: very difficult, 5: very easy) the ease of understanding and implementing the SPARC guidelines for each task.

## Results

Overall, we found that SODA reduced the time required to prepare and share a dataset according to the SPARC guidelines by 70% and made it relatively easier to understand and implement the requirements (Figure 2). After evaluating the shared datasets on Pennsieve, we found that there was an aggregate of 23 compliance errors in datasets submitted without SODA while only one error was found amongst datasets submitted with SODA. When subsequently asked if they would consistently prepare and share their data (SPARC or else) if not mandatory, a majority (8/10 yes, 2/10 maybe) responded that they would if a software application like SODA was available while a majority would not without it (6/10 no, 4/10 maybe). After improving the user flow, adding more automation, and additional integration with other SPARC resources since the beta test, we believe that SODA is reducing researchers’ time, effort, and error in the process even more now.

**Figure 2:**
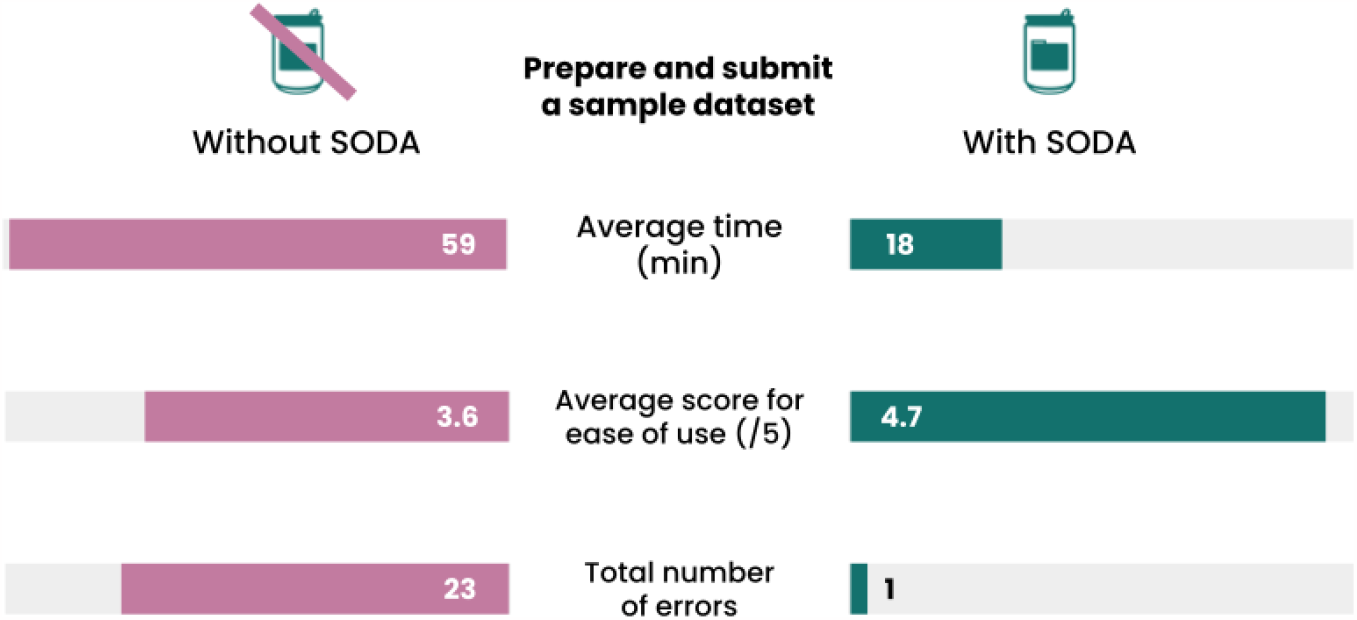
Results from testing of SODA by 10 beta testers during 2020.

Given these advantages offered by SODA, it has been widely used (over 2,000 downloads) by SPARC researchers since 2020 and by researchers collecting ANS-related data outside of SPARC since 2022, when sparc.science became an open repository. Since the beginning of 2021, SODA has helped researchers all over the world process over 16TB of data corresponding to over 250k individual data files.

## Discussion

There is a major push to make data FAIR in all fields of research, including biomedical research with the NIH’s leadership. As a result, many standards, guidelines, and platforms to archive data are developed to achieve that. However, the burden of understanding, learning, and using these resources for making data FAIR is mostly left to the researchers. SODA shows that the availability of an easy-to-use tool can reduce researchers’ time, effort, and error in the process of making data FAIR.

To our knowledge, SODA is the first researcher-oriented tool for making data FAIR. It has since inspired several other tools we are developing such as FAIR-share (Soundarajan and Patel 2023) through support from the National Institute of Allergy and Infectious Diseases (NIAID) and fairhub.io (Soundarajan, Gasimova, and Patel 2023) through support from the NIH Bridge2AI Program. The codebase of SODA was also forked by another team that is developing a tool called NWB GUIDE to simplify the process of preparing and sharing data from the NIH Brain Initiative Program. SODA has itself been made FAIR in line with the FAIR-BioRS guidelines (Patel, Soundarajan, Ménager, et al. 2023) to promote and facilitate such reuse of its source code outside of the developing team.

As the development of SODA continues, we aim to continue improving the user flow, adding more automation, and additional integration with other SPARC resources. We hope that SODA will inspire other biomedical research communities to develop researcher-oriented tools for making data FAIR so that the burden is reduced on researchers and FAIR practices are broadly adopted by them.

## Code Availability

The code associated with this manuscript consists of the source code of SODA and the source code of the SODA documentation. The source code for SODA is hosted on GitHub (https://github.com/fairdataihub/SODA-for-SPARC). It was made FAIR as per the FAIR-BioRS guidelines, and shared on Zenodo (Soundarajan et al. 2023) under the permissible MIT license. The source code for the FAIRshare documentation is maintained on GitHub as well (https://github.com/fairdataihub/SODA-for-SPARC-Docs) and is shared under the permissible MIT license on Zenodo (Patel, Soundarajan, Marroquin, et al. 2023b).

## Acknowledgements

This work was supported by grant NIH SPARC OT2OD030213. We thank the SPARC Data Resource Center (DRC) teams for their continuous support in the development of SODA.

